# Effect of therapeutic ultrasound on the mechanical and biological properties of fibroblasts

**DOI:** 10.1101/2021.11.22.469508

**Authors:** Rosy P. Cárdenas-Sandoval, Homero F. Pastrana-Rendón, Alba G. Ávila-Bernal, Angélica M. Ramírez-Martínez, Myriam L. Navarrete-Jimenez, Alejandro O. Ondo-Mendez, Diego A. Garzón-Alvarado

## Abstract

Conventional doses of therapeutic ultrasound alter the mechanical behavior of ligament fibroblasts to improve the regenerative and remodeling stages of the wound healing process. Using a multidisciplinary approach, we applied ultrasound doses of 1.0 and 2.0 W/cm^2^ at 1 MHz frequency for five days on ligament fibroblasts. Atomic force microscopy showed a decrease in cell elastic modulus for both doses, but the treated cells were still viable based on flow cytometry. Finite element method analysis exhibited visible cytoskeleton displacements and decreased harmonics in treated cells. Colorimetric assay revealed increased cell proliferation, while scratch assay showed increased migration at low doses. An increase in collagen and fibronectin was detected by enzyme-linked immunoassay at high doses, and β-actin expression for both treatments was visualized through immunofluorescence imaging. Both doses of ultrasound altered the fibroblast mechanical properties due to cytoskeletal reorganization and enhanced the early and late stages of cell repair.

## Introduction

Therapeutic ultrasound produces sound waves to create vibrations that exert forces on cells and stimulate the regenerative and remodeling stages coordinated by fibroblasts during the wound healing process [1–3]. Nevertheless, the exact dose of ultrasound that may affect the ligament fibroblast elastic modulus and harmonic vibration to improve the regenerative and remodeling stages remain largely elusive. The elastic modulus of cells is a biomarker that determines several biological responses such as communication with the environment, cell death, aging, and cellular motility [4–6]. It may also influence the harmonic vibration (i.e., the natural frequencies of vibration of the cell structure) [7], which is the rate at which the structure oscillates at a point of balance without being affected by an external force [8].

Therapeutic ultrasound generates a micro-massaging effect caused by compression and negative pressure resulting from micro-vibration and cavitation [9–13]. Cellular transmembrane receptors such as integrins and cadherins detect these forces, and consequently, external stimuli are conducted rapidly along cytoskeleton filaments and absorbed at reserved edges in the cytoplasm and nucleus, modifying the cellular genome activities by increasing collagen synthesis and activating the mitotic activity of cells [14–16].

Cells react to external physical stimuli caused by ultrasound by altering their cytoskeleton, which is the structure responsible for regulating the mechanical behavior of cells. The cytoskeleton maintains the cell shape, responds to external mechanical cues, exerts forces, and produces motion [13,17,18]. It transduces the mechanical signal and converts it into a biological response associated with the wound healing process. This is evidenced by measuring changes in the cell mechanical properties [4–6,17,19–22].

The mechanical and biological effects of applying conventional ultrasound doses of 0.1–3 W/cm^2^ spatial average temporal average intensity (SATA) at 1–3 MHz frequency to ligament fibroblasts remains unclear [23]. Evidence so far has shown contradictory results: on the one hand, when therapeutic ultrasound is applied at conventional frequencies [9–12], there is no resonance effect. For example, when applying therapeutic ultrasound at 1 MHz and low intensities (less than 1.0 W/cm^2^), cell proliferation and extracellular matrix (ECM) improve on dermal and periodontal ligament fibroblasts [23–27]. Moreover, this high frequency may modulate cell cytoskeleton organization [28, 29] which in turn stimulates cell migration [30]. On the other hand, when applying ultrasound at low frequencies in the kilohertz range, a resonance effect occurs, in which the integrity of the cell structure may be compromised because of the oscillation at such frequencies that coincide with the harmonic vibration of the cell [7,31–33]. For example, when applying ultrasound at frequencies between 550 kHz and 650 kHz, cell death increases and cell proliferation decreases in breast cancer cells [34].

The effects of conventional doses of therapeutic ultrasound used in rehabilitation [23] on the elastic modulus and harmonic vibration (mechanical properties), as well as on the viability, proliferation, migration, and synthesis of the ECM (type I collagen, type III collagen, and fibronectin) and β- actin expression (biological properties) of ligament fibroblasts in joints remain uncertain [35–38]. The controversy concerning the contradictory results of ultrasound and the lack of evidence motivated us to measure and demonstrate that the mechanical and biological properties can be modulated by the reorganization of the ligament fibroblast cytoskeleton after applying conventional doses of therapeutic ultrasound.

Therapeutic ultrasound modifies the cell elastic modulus and its harmonic vibration, while cells reorganize their cytoskeleton structure without resonance or harmful effects. It also improves the regenerative and remodeling functions of ligament fibroblasts. This complex effect cannot be evaluated by clinical assessment of a physiotherapist. Thus, we applied a multidisciplinary approach that integrates *in vitro* and computational techniques to assess the effects on the mechanical and biological responses of ligament fibroblasts of joints by applying two intensities of conventional therapeutic ultrasound, namely, a low dose of 1.0 W/cm^2^ and a high dose of 2.0 W/cm^2^, both at a frequency of 1 MHz. Furthermore, we determined the specific ultrasound dose required to improve the regenerative (early) and remodeling (late) phases of the ligament fibroblast healing process.

## Materials and Methods

An explant technique was used to obtain ligament fibroblasts. The protocol of ultrasound therapy applied to cultured cells followed the standard clinical guidelines for ligament treatment. We evaluated the effects of therapeutic ultrasound on the mechanical properties of ligament fibroblasts by measuring their elastic modulus using atomic force microscopy (AFM), and the harmonics of the natural frequencies of vibration of the ligament fibroblast cytoskeleton using a finite element method FEM analysis. Moreover, we evaluated the biological properties of ligament fibroblasts by performing the following experiments: cell viability using flow cytometry, fibroblast cell proliferation by a colorimetric assay), cell migration using scratch assay, ECM synthesis by enzyme-linked immunoassay (ELISA), and β-actin expression through immunofluorescence imaging. A diagram of the methodology is shown in Fig 1.

**Fig 1.**
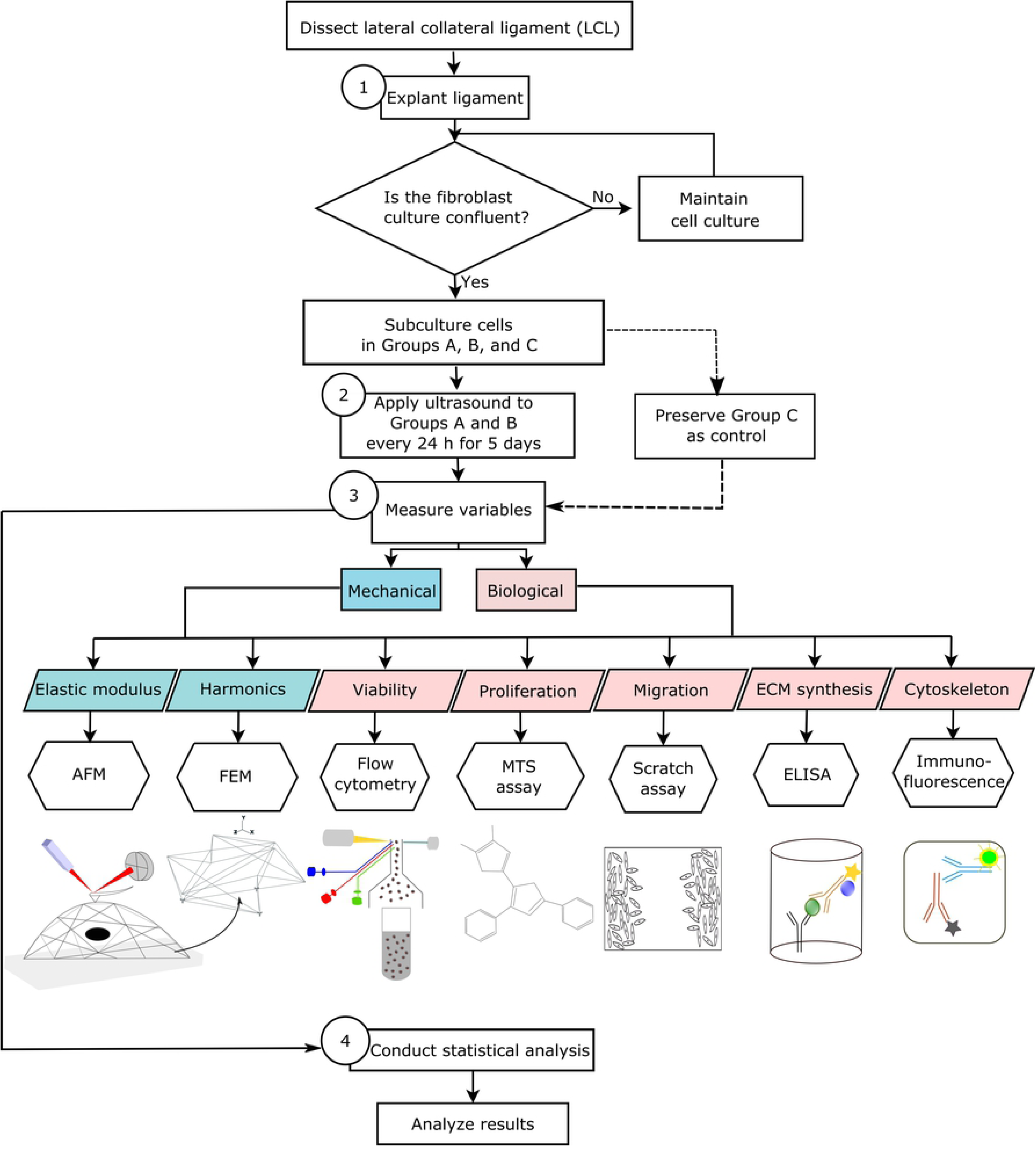
Diagram of the methodology. (1) Ligament explant, (2) ultrasound stimulation, (3) measurement of mechanical and biological parameters, and (4) statistical analysis.

### Ligament fibroblast explant

Fibroblast cells were obtained using an explant technique from the lateral collateral ligaments (LCLs) of both knees of one adult male Wistar rat, as depicted in Fig 2A–B, following a previously described protocol [39]. The experiments were performed according to the international regulations of laboratory animals. Moreover, the animal studies and research protocols were approved by a local ethics committee (Protocol Number:FC-13-01082016). The LCLs were maintained under aseptic conditions in a 1:1 mixture of Dulbecco’s modified Eagle medium and nutrient mixture F-12 (DMEM/F12; DF-041-B; Merck Millipore) supplemented with 1% antibiotic/antifungal (15240062; Gibco) and 10% fetal bovine serum (FBS; 12657029; Gibco). Dissection was performed with the aid of a scalpel to cut the femoral and fibular insertions. The LCLs were placed into T-25 culture flasks containing a sterile supplemented culture medium, as exhibited in Fig 2C. The culture medium was changed every 48 h.

**Fig 2.**
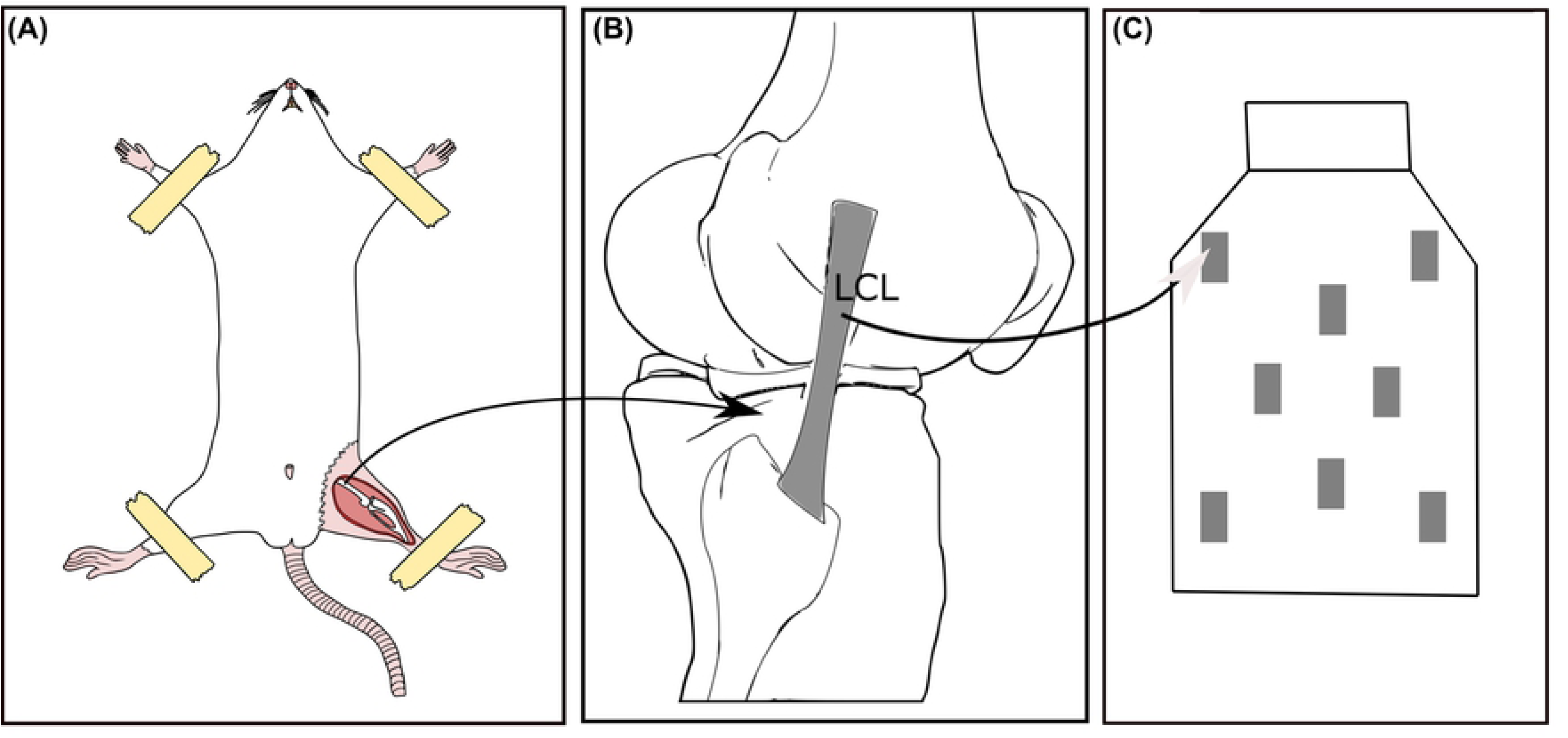
Ligament fibroblast explant. (A) Adult male Wistar rat; (B) lateral collateral ligament (LCL); (C) ligament tissues cultured in a T-25 culture flask.

The flasks were incubated at 37°C in a humidified atmosphere containing 5% CO_2_. After 15 days, the monolayer cultures became confluent, and the tissues were removed from the flasks. Explanted cells were washed with Hank’s balanced salt solution (14065056; Gibco), detached using 0.025% trypsin (15400054; Gibco) for 5 min, centrifuged at 287×g for 5 min, and subcultured for subsequent experiments. The remaining cells were cryopreserved in a mixture of 10% DMEM/F12, 80% FBS, and 10% dimethyl sulfoxide.

To ensure that the cells explanted from the ligaments corresponded to fibroblasts, their nuclei and bodies were highlighted using hematoxylin-eosin staining (H&E) and microscopic observation [40]. We followed the laboratory protocol described by Spitalnik [41].

### Pulsed ultrasound intensity and application time

We applied energies of 1.5 and 5.0 J/cm^2^ for five days to the two treatment groups that we designed. These energies correspond to 5% and 17% of the 30 J/cm^2^ applied in human therapy [42–46]. Treatment Group A received 1.0 W/cm^2^ (low dose), whereas treatment Group B received 2.0 W/cm^2^ (high dose) of the maximum intensity. The SATA values were 0.5 W/cm^2^ and 1.0 W/cm^2^ for treatment Groups A and B, respectively. A layer of sound wave transmission gel was applied over the transducer, and the culture plates were then placed over the ultrasound device. Control cells were processed in the same way but without turning the device on. For both treatments, a standard “J-Style” (JC-2902) ultrasound device was used following the standard clinical procedures for ligament therapy [47].

Based on the ultrasound parameters and the surface area of the plate culture, the application time *t* of the ultrasound was calculated using the potency equation of energy transmission [11,12,48], expressed as Equation 1.

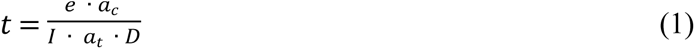

Here, *e* is the energy per square centimeter [49], *I* is the transducer intensity, *a_t_* is the effective radiated area, *D* is the duty cycle, and *a_c_* is the culture area. The application time for each culture plate is listed in Table 1. The control group used the same surface area of plate cultures for every experiment as that in the treatment groups with no ultrasound.

**Table 1.** Surface area and application time of ultrasound in each treatment group.

| Experiment variable | Surface area of plate culture (cm<sup>2</sup>) | Ultrasound application time (s) |  |
| --- | --- | --- | --- |
|  |  | Group A (low dose: 1.0 W/cm<sup>2</sup>) | Group B (high dose: 2.0 W/cm<sup>2</sup>) |
| *Elastic modulus*<br>Petri dish | 11.78 | 7 | 12 |
| *Viability*<br>6-well plate | 3.5 | 2.1 | 3.5 |
| *Cell proliferation*<br>96-well plate<br>(groups of 9 wells) | 5 | 3 | 5 |
| *Migration*<br>12-well plate | 109.9 | 66 | 110 |
| *ECM synthesis*<br>6-well plate | 3.5 | 2.1 | 3.5 |

### Fibroblast elastic modulus by AFM

Fibroblasts (3.5 × 10^2^ cells from each treatment group) were cultured in Petri dishes (3.5 cm diameter) and reached 20–40% confluency on the 6^th^ day. We measured the elastic modulus of proliferating viable cells maintained in DMEM/F12 within 2–3 h. Changes in the cell elastic modulus were monitored using AFM (MFP3D-Bio AFM system, Asylum Research, Santa Barbara, CA) [50, 51].

Soft cantilevers T R400P B (Olympus, Japan) with a nominal spring constant of 0.09 N/m, a tip radius of 42 nm, and a half-opening angle of 35° were employed. The relative trigger force was 2 nN. The elastic modulus was estimated as a function of the position on the cell using the force–volume technique by measuring the cantilever deflection. We used a video microscope to position the AFM tip at precise locations over the cell surface. The probe moved up and down, simultaneously registering the force curve and cell topography at each pixel of the surface. We obtained force–volume images with a resolution of 20 × 20 pixels within 30 × 30 µm^2^ scan areas for 10 cells per group. The approximate acquisition time per image was 15 min.

The numbers of effective measurements performed for Groups A, B, and C were 863, 866, and 338 indentations, respectively. We used a larger sample to better calculate the average values of the elastic modulus and avoid errors associated with the indentation depth. Force curves determined from a relative area above the whole cell enabled the comparison of induced changes to a constant force. Because the elastic modulus ( calculated using the Sneddon model) may create a substrate effect, which is a source of error, due to sharp probe indentation producing a larger depth [52], we used an asymptotical correction model to nullify the bottom substrate effect for rigid cone indenting [53], as indicated in Equation 2.

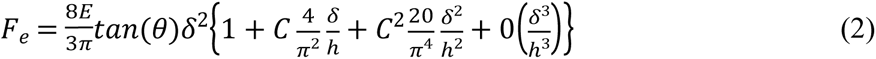

Here, *F_e_* is the elastic force, *E* is the elastic modulus, *δ* is the indentation depth, θ is the rigid cone angle (set as 35°), *h* is the thickness of adherent cells at the point of indentation (set as 150 nm), 0 represents higher-order terms in the series (assumed to be negligible), and *C* = 1.7795 tan(*θ*) [53]. Refer to the supporting information (S1 File) for additional details.

### Harmonic vibration and modal analysis by finite element method (FEM)

We proposed a simplified theoretical model to consider only the cytoskeleton, as it predominantly determines cell mechanics and its response to external stimuli. Without the network of filaments that compose the cytoskeleton (actin, intermediate filaments, and microtubules), fibroblasts will be extremely deformable to preserve their shape and biological response (proliferation, migration, and synthesis of the ECM) in the wound healing process [54, 55]. Then, we performed an FEM analysis to simulate the harmonics of the natural frequencies of vibration of the ligament fibroblast cytoskeleton using the eigenvalue extraction method Lanczos (ABAQUS/CAE 6.12.3 software). Modal analysis was used to predict the 50^th^ natural frequencies and eigenforms of the cytoskeleton. A three-dimensional (3D) octahedron tensegrity model with 12 coordinates and 30 beam elements represented the cytoskeleton (Fig 3) [56]. As the cytoskeleton is composed of a protein network of filaments, the tensegrity structure mimics 12 actin filament beams, 12 intermediate filament beams, and microtubule beams. The actin filaments were located in the cortex, the intermediate filaments in the middle, and the microtubules were in the center of the structure [57, 58].

**Fig 3.**
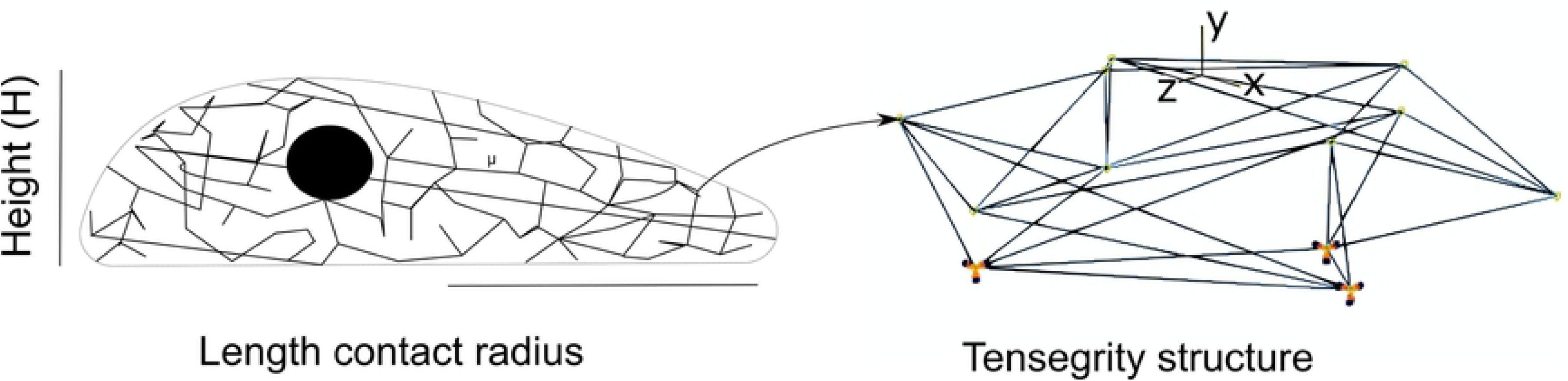
Three-dimensional octahedron tensegrity model. Ligament fibroblast cytoskeleton tensegrity structure.

The cell elastic modulus measured by AFM may elucidate the changes in cytoskeleton elasticity [50, 59]. Hence, the elastic modulus median and the mean height values for treated cells and the control group were taken from the AFM results as input parameters for the tensegrity structure to calculate the harmonic vibration and perform the modal analysis for every cytoskeleton configuration in each group. The data for the Poisson’s ratio, length, and beam radius of the tensegrity structure were taken from the literature. These values are provided in Table 2 [60–63].

**Table 2.**
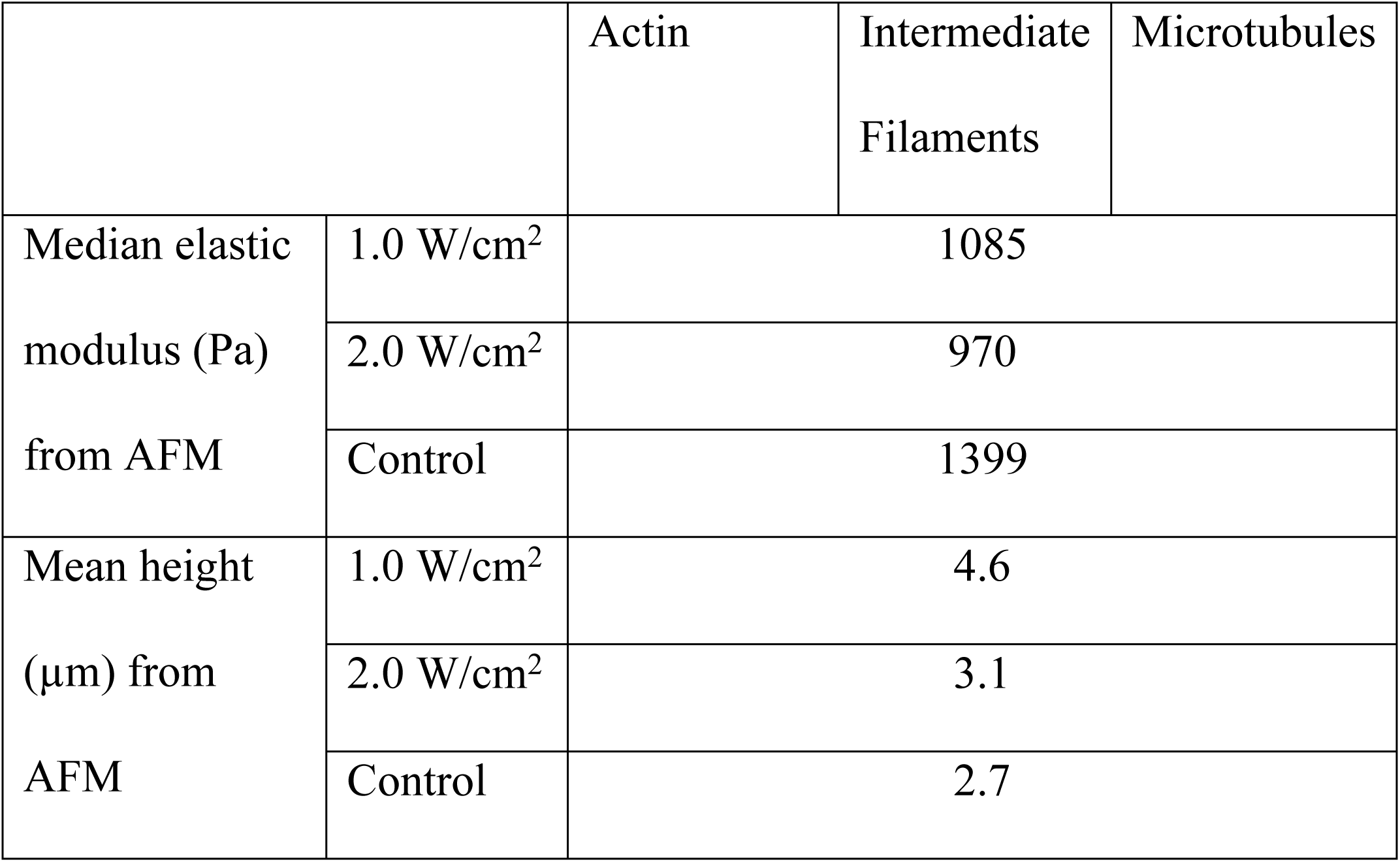
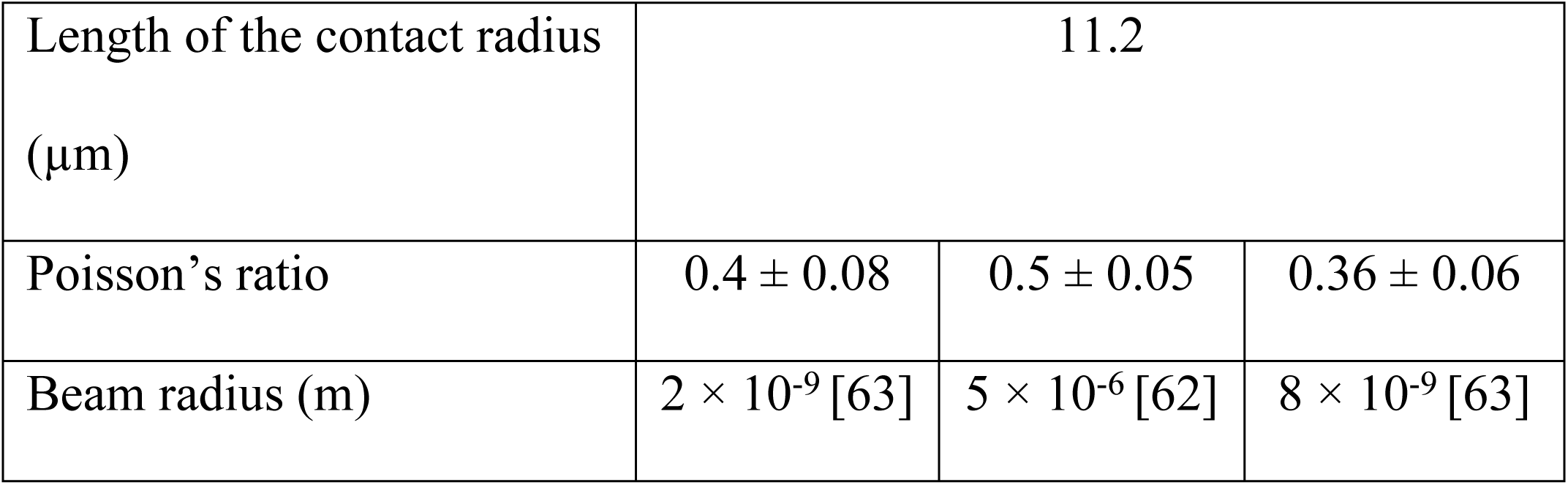
Cytoskeleton filament material characteristics. The elastic modulus and height were taken from the AFM results obtained for both treatment groups and the control group. The Poisson’s ratio, length, and beam radius were taken from the literature.

|  |  | Actin | Intermediate<br>Filaments | Microtubules |
| --- | --- | --- | --- | --- |
| Median elastic<br>modulus (Pa)<br>from AFM | 1.0 W/cm<sup>2</sup> | 1085 |  |  |
|  | 2.0 W/cm<sup>2</sup> | 970 |  |  |
|  | Control | 1399 |  |  |
| Mean height<br>(μm) from<br>AFM | 1.0 W/cm<sup>2</sup> | 4.6 |  |  |
|  | 2.0 W/cm<sup>2</sup> | 3.1 |  |  |
|  | Control | 2.7 |  |  |

|  |  |  |  |
| --- | --- | --- | --- |
| Length of the contact radius<br>($\mu\text{m}$) | 11.2 | | |
| Poisson's ratio | $0.4 \pm 0.08$ | $0.5 \pm 0.05$ | $0.36 \pm 0.06$ |
| Beam radius (m) | $2 \times 10^{-9}$ [63] | $5 \times 10^{-6}$ [62] | $8 \times 10^{-9}$ [63] |

The units were converted to microscale. The cytoskeleton filaments were considered isotropic and elastic because the cells had small deformations of 2–8% [57,63–65]. The beam length of the contact radius in the tensegrity structure was 11.2 µm [63]. The filament density was 1.15 × 10^-6^ µg/µm^3^ [7]. The initial boundary conditions were imposed on the three base nodes of the tensegrity structure [63]. The three receptor nodes represented the focal adhesion of the cell to the ECM because ligament fibroblasts are adherent cells and are dependent on the actin cytoskeleton [66]. They were constrained for three degrees of freedom (U1 = U2 = U3 = UR1 = UR2 = UR3 = 0).

The height values (Y axes) for each octahedron structure were taken from the mean values obtained in the AFM force–volume topography maps, and the X and Z axes values were taken from the literature to comply with the spread shape of an adherent cell [63, 67].

### Fibroblast viability measured using flow cytometry

Fluorescein isothiocyanate (FITC)-conjugated Annexin-V (ab14085) and cationic marker propidium iodide (PI-ab14083) were used to quantitate non-apoptotic cells, cells in early apoptosis, necrotic cells, and cells in late apoptosis post-treatment [68]. Samples were analyzed using a flow cytometer (BD FACS Canto II) with a solid-state (L1) laser (488 nm excitation line, air-cooled, 20 mW solid state).

Fibroblasts (2.84 × 10^4^ cells per group) were cultured in a six-well plate until 100% confluency was attained on the 6^th^ day. One well was stained with Annexin V-FITC, another with PI, and the third with both. The other wells served as controls. The samples were placed in cytometry tubes and diluted in 300 μL of the culture medium. Each well corresponded to a specific sample. Ten thousand events were recorded to identify non-apoptotic cells (Q3: Annexin V-FITC negative/PI negative), early apoptotic cells (Q4: Annexin V-FITC positive/PI negative), necrotic cells (Q1: Annexin V-FITC negative/PI positive), and late apoptotic cells (Q2: Annexin V-FITC positive/PI positive) [69].

### Fibroblast cell proliferation by MTS assay

An MTS tetrazolium assay (MTS=3-(4,5-dimethylthiazol-2-yl)-5-(3-carboxymethoxyphenyl)-2-(4-sulfophenil)-2H-tetrazolium, inner salt) [70] measures the interaction between viable cells (mitochondrial enzymatic activity) and phenazine methosulfate. This interaction generates a formazan product that is soluble in culture media to count the increased number of viable fibroblast cells [71–73]. We quantified the number of viable fibroblast cells in proliferation on the 1^st^ and 3^rd^ day of the stimulation period and on the 6^th^ day after the stimulation period.

Ligament fibroblasts from each treatment group were cultured in a single 96-well plate (4 × 10^1^ cells/well) containing 200 µL of culture medium until 100% confluency was achieved on the 6^th^ day. We divided the 96-well plate into six groups of nine wells (3 × 3 wells). The remaining wells served as blank controls, in which only a culture medium with no cells was added. Twenty microliters of reagent was added directly to each well plate following the instructions of the MTS Cell Proliferation Assay Kit (colorimetric ab197010, abcam®). The culture plate was incubated for 3 h at 37°C in a humidified atmosphere containing 5% CO_2_. Absorbance at 490 nm was measured using a microplate reader. The mean absorbance was calculated by subtracting the absorbance of the treatment groups from the absorbance of the blank controls.

### Fibroblast cell migration measured by scratch assay

We seeded 50 × 10^3^ cells in a 12-well plate to fully cover the surface after finishing the treatment (6^th^ day). We used an *in vitro* scratch assay as described by Liang et al. to measure ligament fibroblast migration [74]. The culture medium lacking FBS was changed post-treatment to avoid cell division. Each well of the plate was scratched using a 10 µL tip. Images at 0 and 24 h (when fibroblast migration closed the scratch for each group) of post-treatment were captured using a Cytation 3 Cell Imager Multi-Mode Reader (Biotek) and processed using Gen 5.2.0.7 software (Biotek).

The measurements of migration were single-blinded. Six migration measurements were registered with respect to a vertical line located at the center of the image along with each cell distance (1 pixel equivalent to 1 µm). We measured the distance that every cell moved (mean value) by calculating the difference between two measurements. Next, we compared the beginning (0 h) to the end of the 24 h period of migration using ImageJ software version 1.50i 3 for Windows (developed by Wayne Rasband, National Institutes of Health, USA, http://imagej.nih.gov/ij).

### Fibroblast ECM synthesis: type I collagen, type III collagen, and fibronectin measured by ELISA

Ligament fibroblasts from both treatment groups and the control group were cultured in three separate six-well plates (1.8 × 10^3^ cells/well) containing 1 mL of culture medium until 100% confluency was reached on the 6^th^ day. Only three of the six wells were seeded in the third plate where control Group C was cultured. We used ELISA to measure the protein concentration of type I collagen, type III collagen, and fibronectin in ligament fibroblast supernatants on the 6^th^ and 10^th^ day after the stimulation period [75, 76]. On the 10^th^ day, we evaluated the protein concentration because type I collagen requires more days to be released than type III collagen. Between the 6^th^ and the 10^th^ day, the cells were maintained under aseptic conditions in a 1:1 mixture of DMEM/F12 (DF-041-B; Merck Millipore) supplemented with 1% antibiotic/antifungal (15240062; Gibco) and 10% FBS (12657029; Gibco). All cell culture supernatant samples (1 mL) were collected on the 6^th^ and 10^th^ day, then stored at 4°C and assayed within 7 days.

We used rat collagen type I (E-EL-R0233), rat collagen type III (E-EL-R0235), and rat fibronectin (E-EL-R0578) Elabscience® ELISA kits. Eight serial dilutions were performed using the reference standard from the kit until the protein reached 20 ng/mL. Then, we collected a duplicate sample of each dilution, and 16 samples were transferred to a 96-well ELISA plate. The culture medium (100 μL/well) from both treatment (6 samples) and control (3 samples) groups on the 6^th^ and 10^th^ day was aspirated and transferred to a 96-well ELISA plate (a total of 30 samples). We added a duplicate blank control (100 μL/well of the standard working solution of the substratum) and a duplicate negative control (100 μL/well of the supplemented cell culture medium and no cells) to the 96-well ELISA plate.

The readings of the wells that contained the samples of the treatment and control groups, blanks, and negatives were obtained using a microplate reader (absorbance of 450 nm). The reading of the negative control was subtracted from the readings of the treatment and control groups to obtain the final optical density (OD) of the samples. A curvilinear regression line was constructed using the standard to calculate the protein concentration. In total, six samples from the treatment groups and three from the control group were measured. When the concentrations were less than the lowest value of the standard curve, the protein concentration (ng/mL) was not detected in one sample from treatment Group A on the 10^th^ day, in three samples of treatment Group B on the 6^th^ day, and in two samples of control Group C on the 6^th^ day.

### β-actin expression measured by immunofluorescence imaging

To demonstrate that ligament fibroblasts activate their cytoskeleton after treatment, we assessed the presence of β-actin filaments, an essential component of the fibroblast cytoskeleton [77]. Treated ligament fibroblasts cultured on coverslips were fixed with 4% formaldehyde, permeabilized using 0.1% Triton, stained with a monoclonal antibody (Invitrogen,15G5A11/E2) against rat β-actin at a dilution of 1:2000, and incubated overnight at 4°C. Afterward, the cells were incubated with 488-conjugated goat anti-mouse IgG secondary antibody (Invitrogen, A32723) at a dilution of 1:1000 conjugated with Invitrogen Alexa Fluor®. Finally, nuclei were stained with Hoechst 33342. Images were taken on a cell imaging reader microscopy (Cytation 3 cell imager multi-mode reader; Biotek).

We calculated the shortening area of β-actin in each image corresponding to each treatment group. Then, we analyzed three images per group using the immunohistochemistry (IHC) image analysis toolbox plugin in ImageJ software version 1.50i 3 for Windows (developed by Wayne Rasband, National Institutes of Health, USA, http://imagej.nih.gov/ij) by following the live-cell actin analysis proposed by Hoyle et al. [66]. We trained the algorithm by recording five color pixels of the images to detect the green color of the β-actin filaments stained with Alexa Fluor®. The statistical color detection model was then automatically calculated. A total of 29 ± 1 cells per image were analyzed per group. After generating the detected green area in each image, we calculated the total area of the β-actin filaments using the following steps. First, we converted the read-green-blue (RGB) image into an 8-bit grayscale. Then, we adjusted the threshold to select the entire β-actin green area and converted it to black. (Additional details can be found in the supporting information (S2 File)). Finally, we run the particle analysis by defining the size (zero to infinity) and circularity (0.0 to 1.0). The results were obtained in micrometers.

### Statistical analyses

Statistical analyses were conducted using BioVinci software version 2.8.5 for Windows (BioTuring Inc., San Diego California USA, www.bioturing.com), a software environment for data visualization, analysis, and machine learning in the life sciences. All data were representatives of at least three independent experiments. Normality was assessed using the Shapiro–Wilk test, Pearson chi-square test, one-sample Kolmogorov–Smirnov test, and Jarque–Bera test. Because the data were not normally distributed, the medians (elastic modulus and ECM synthesis) and means (fibroblast cell proliferation and migration) were compared across groups using a nonparametric multiple comparison Kruskal–Wallis test. The data were presented as means and standard errors of the means (SEMs) for the cell proliferation, cell migration, and β-actin area. Then, they were shown as a boxplot with medians for elastic modulus, harmonics, and ECM synthesis. Statistical significance was assumed at P < 0.05.

## Results

### Ligament fibroblasts

The ligament fibroblasts were motile around the explanted tissue (Fig 4A) and then adhered to the flasks where they proliferated. The typical characteristics of fibroblasts (Fig 4B), namely, adherence; presence of nucleus and body; flat, elongated, and triangular shape; and linkage between cells, were evident [78].

**Fig 4.**
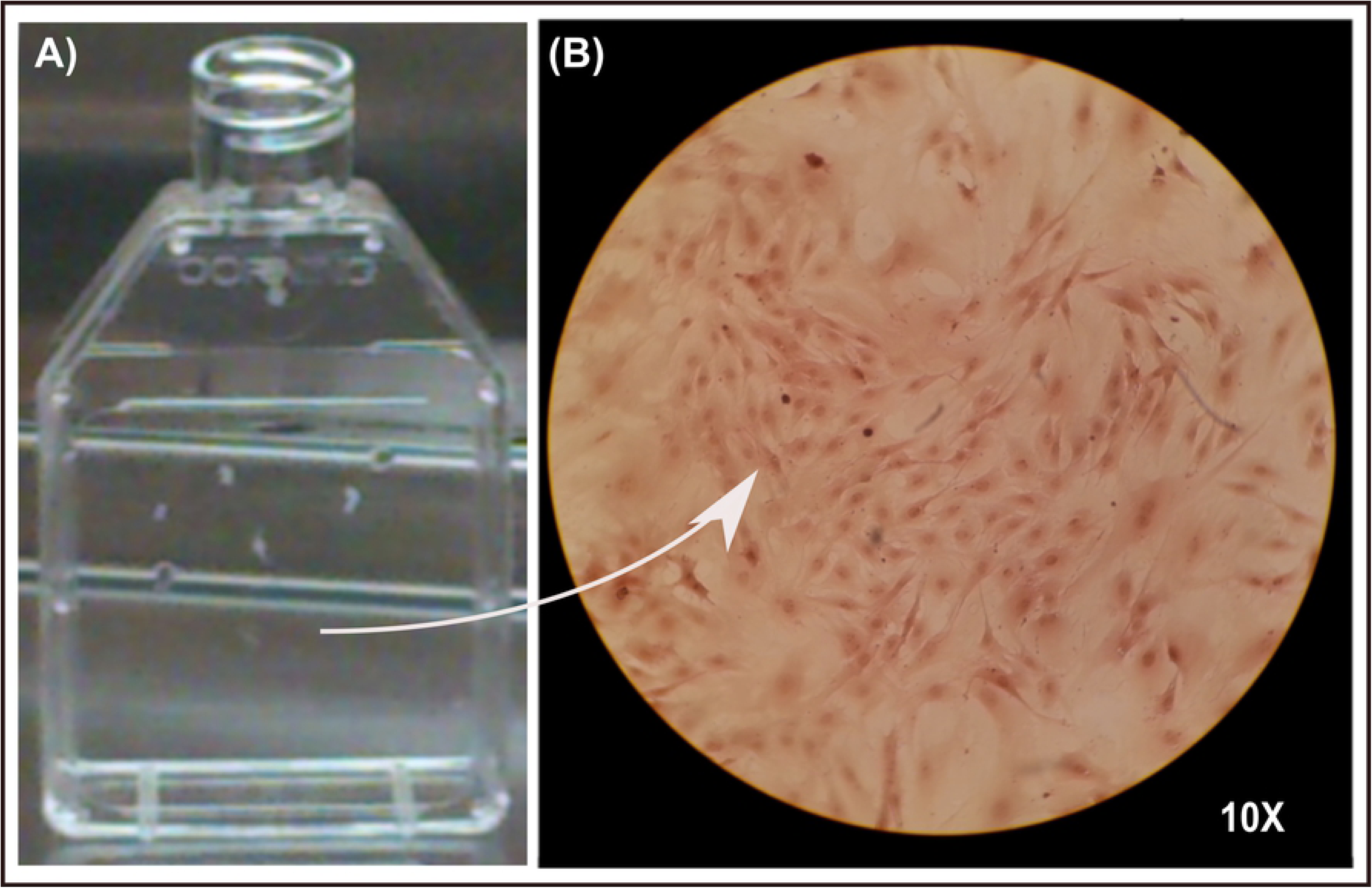
Ligament fibroblasts. (A) After 15 days, the ligament fibroblasts were motile around the ligament explant. (B) Adherent cells were stained with H&E.

### Effect of ultrasound treatment on ligament fibroblast structure

The AFM revealed that the median elastic modulus of the treated ligament fibroblasts decreased by 22% for the low dose (1.0 W/cm^2^) and 31% for the high dose (2.0 W/cm^2^) compared to that of the control group. We found significant differences among the groups (***P = 0.00001 × 10^-6^; Fig 5A). Additionally, the ligament fibroblast topography through 3D force–volume maps showed darker areas for the treated cells than those of the control group (Fig 5B). This means that the application of low and high doses of therapeutic ultrasound softens the ligament fibroblast structure. Because the cell structure is mainly provided by the cytoskeleton, we inferred that both treatments caused a reorganization of the cytoskeleton through β-actin activation, which could produce cell biological responses such as proliferation, migration, and ECM synthesis, as will be demonstrated in the next sections.

**Fig 5.**
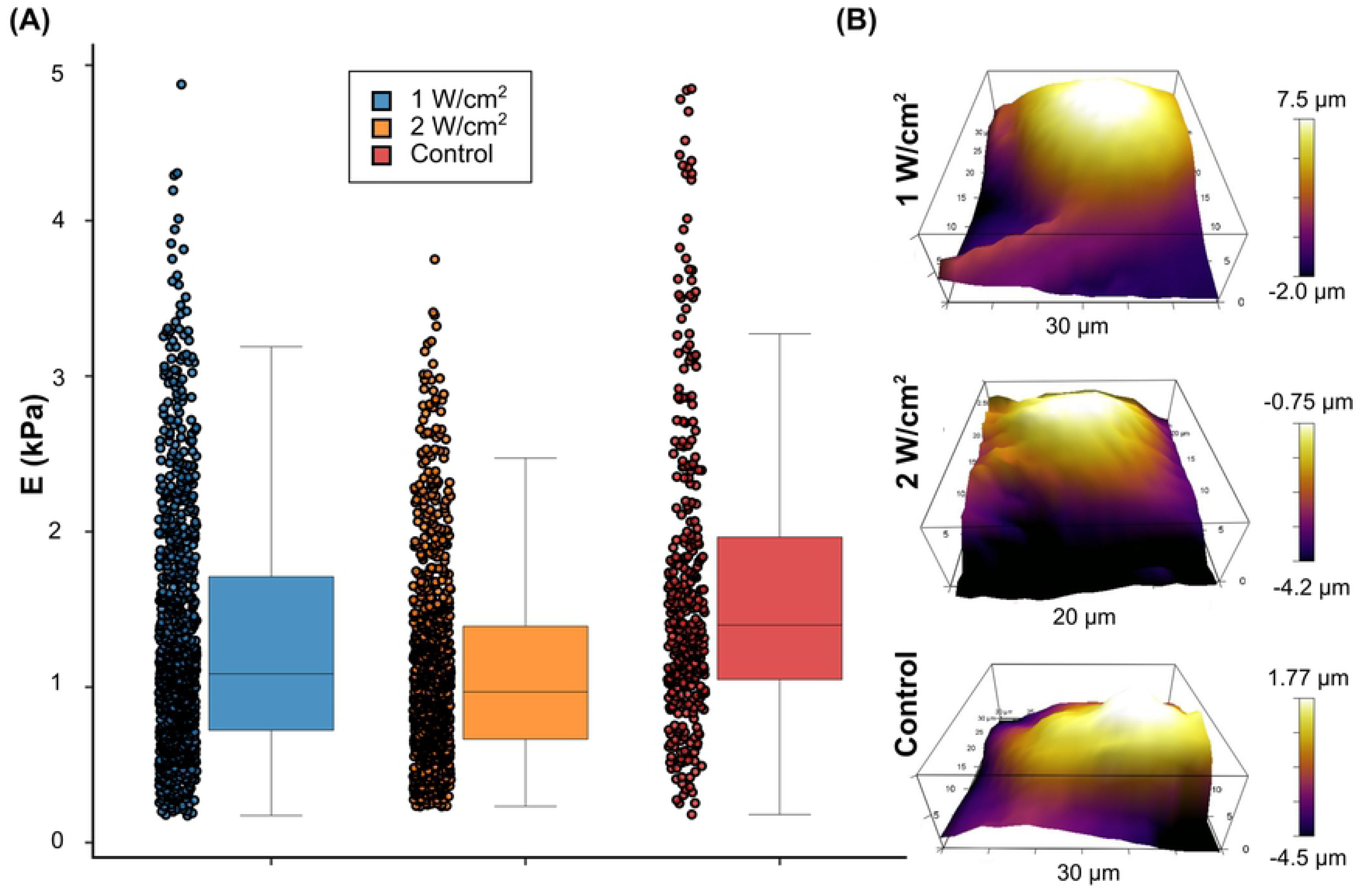
Softening of ligament fibroblast structure due to low and high doses of ultrasound. (A) Boxplot showing medians with whiskers from minimum to maximum values. (B) 3D force– volume topography maps from AFM results.

### Effect of ultrasound treatment on harmonic vibration

Using the FEM, we calculated until the 50^th^ harmonic of the natural frequency of vibration and the 50^th^ eigenform (mode of vibration) for each cytoskeleton structure. We found similar displacements of the ligament fibroblast cytoskeleton for the treated cells but a different displacement for the control group. An example of this finding is illustrated in the 5^th^ mode of vibration (Fig 6A). The displacements for all groups are featured in the animation (S3 Movie). In terms of the natural frequencies of vibration, higher frequencies were predicted when the cytoskeleton filaments had a higher elastic modulus (control group), reaching a maximum vibration frequency of 4.1 × 10^9^ Hz in the 50^th^ vibration mode. When the cytoskeleton filaments had the elastic modulus of treatment Groups A and B from the AFM results, the vibration frequencies decreased up to a maximum of 3.3 × 10^9^ Hz in the 50^th^ vibration mode for both structures (Fig 6B). The median of the 50^th^ harmonic of the natural frequencies of vibration of the treated ligament fibroblasts decreased by 13% for the low dose (1.0 W/cm^2^) and 17% for the high dose (2.0 W/cm^2^) compared to that of the control group. Nevertheless, there was no statistically significant difference among the groups (P = 0.514). It means that harmonic of the natural frequencies of vibration are not dependent on the elastic modulus decreased by the dosage of ultrasound treatment.

**Fig 6.**
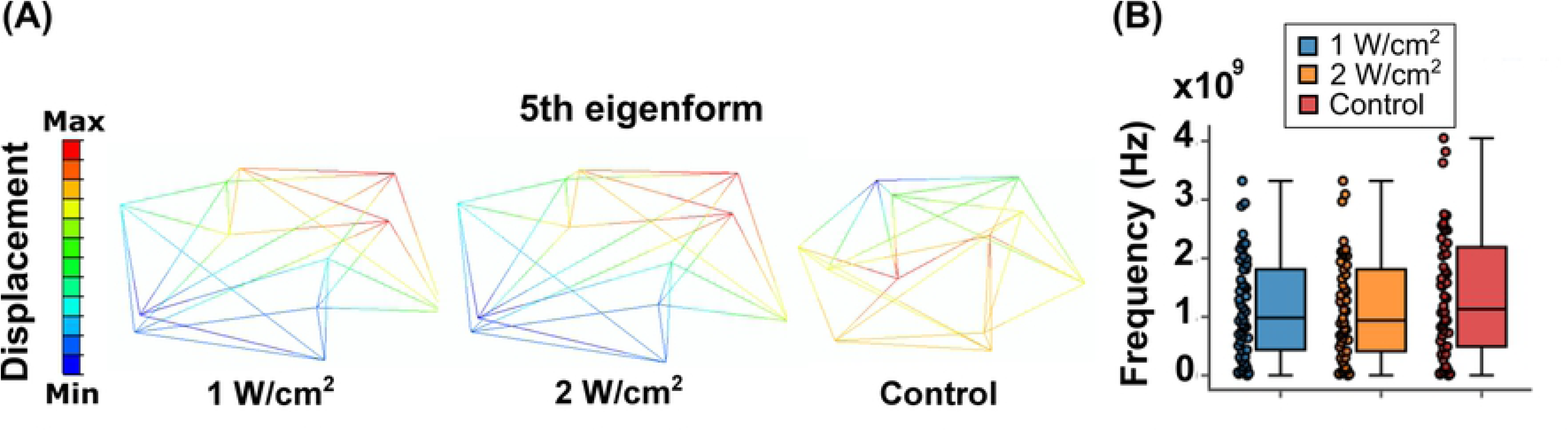
Alteration of ligament fibroblast cytoskeleton due to ultrasound treatment. (A) 3D tensegrity structure for ligament fibroblast cytoskeleton. (B) Boxplot showing medians with whiskers from minimum to maximum values.

### Effect of ultrasound treatment on viability of ligament fibroblasts

Histograms from the cell death assays indicated the number of cells stained with Annexin V-FITC and/or PI (Fig 7A). Flow cytometry showed that most cells were viable in the treatment and control groups (Fig 7B). The number of events collected was 10,000. The results showed that cell viability slightly decreased by 1% for the low dose (1.0 W/cm^2^) and 10% for the high dose (2.0 W/cm^2^) compared to that of the control group.

**Fig 7.**
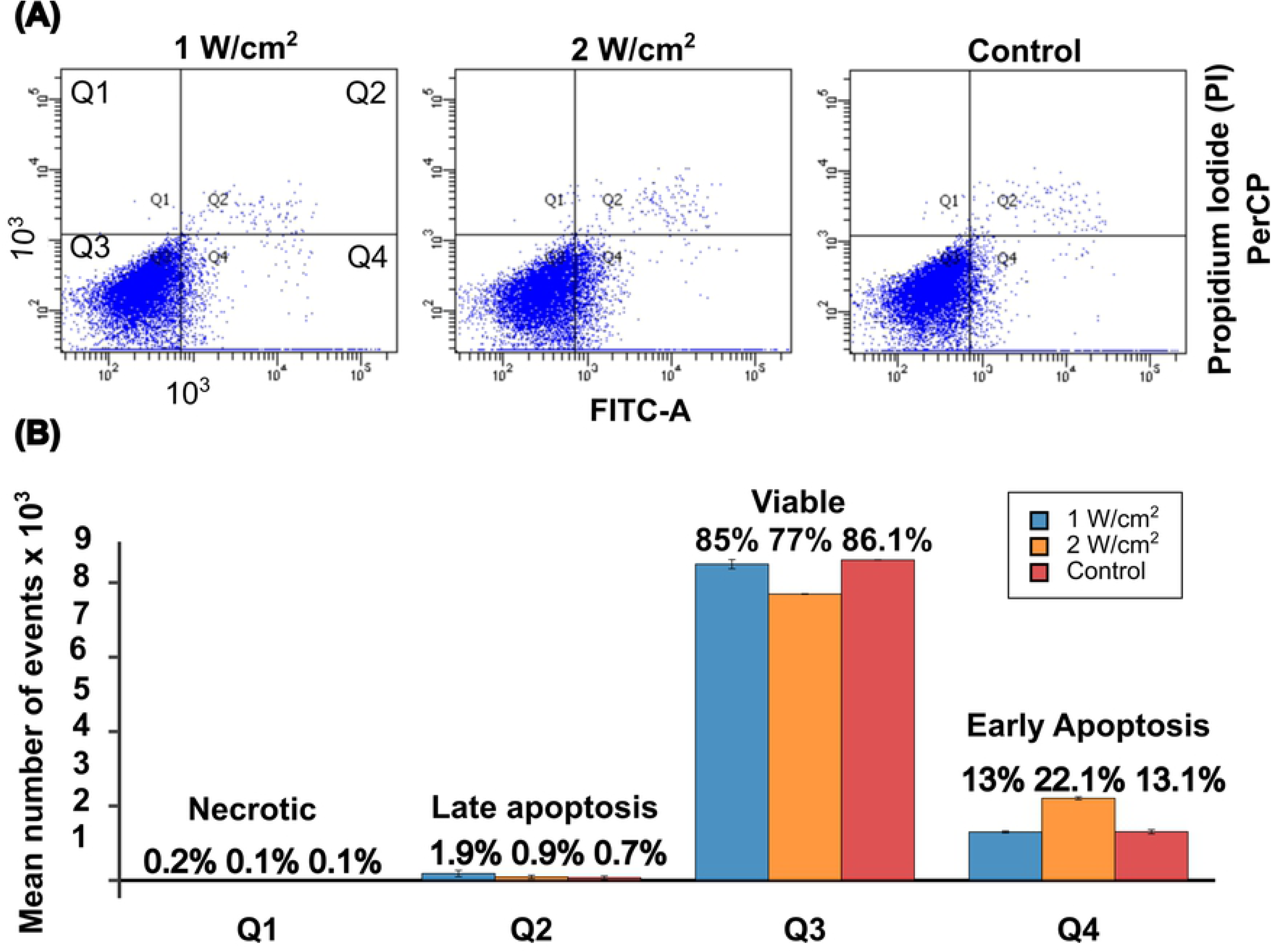
Negligible effect of low and high doses of ultrasound on viability of ligament fibroblasts. (A) Histogram and dot plot of cell viability assay using Annexin V-FITC and cationic marker PI. Quartile 1: necrotic cells; Q2: late apoptotic cells; Q3: viable cells; and Q4: early apoptotic cells. (B) Bar plot of the mean number of events for each quartile of the flow cytometry data.

### Effect of ultrasound on fibroblast cell proliferation in early treatment

The MTS assay showed that on the 1^st^ day of culture, cell proliferation increased by 25% for the low dose (1.0 W/cm^2^) compared to that of the control group. On the 3^rd^ day of culture, the same experiment showed that cell proliferation decreased by 11% for the low dose and 9% for the high dose (2.0 W/cm^2^) compared to that of the control group. On the 6^th^ day of culture (1^st^ post-treatment day), cell proliferation increased by 10% for the low dose but decreased by 13% for the high dose compared to that of the control group. We found significant differences among the groups (*P = 0.041; Fig 8). We demonstrated that a low ultrasound dose increases the cell proliferation of ligament fibroblasts on the 6^th^ day after the stimulation period.

**Fig 8.**
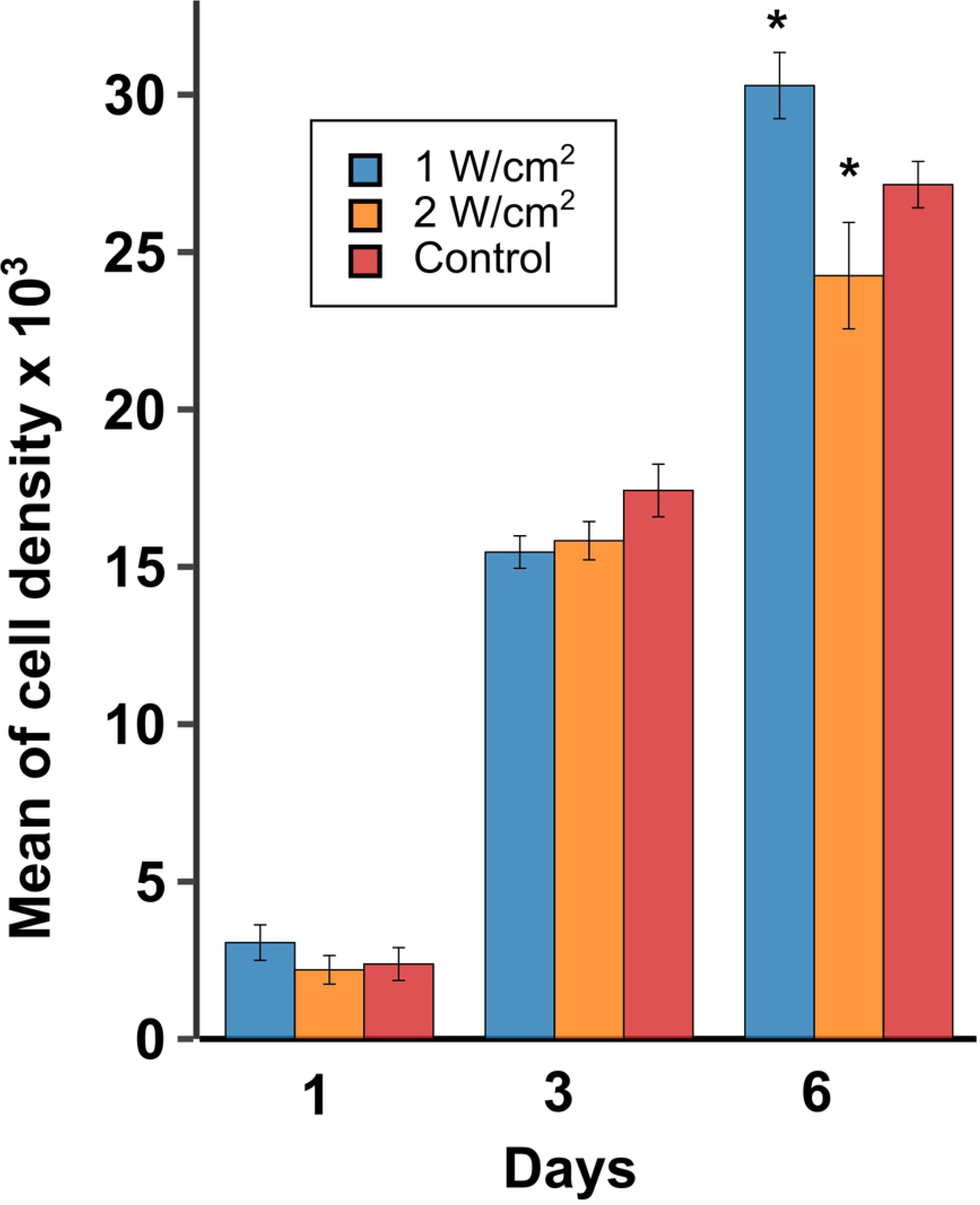
Increase in fibroblast cell proliferation in early treatment due to low dose of ultrasound. The mean of cell number in proliferation of ligament fibroblasts is higher for the low dose on the 6^th^ day after treatment. Error bars indicate SEMs.

### Effect of therapeutic ultrasound on fibroblast cell migration in early treatment

The scratch assay (Fig 9A) showed that the mean migration length of ligament fibroblasts increased by 4% for the low dose (1.0 W/cm^2^) and decreased by 11% for the high dose (2.0 W/cm^2^) compared to that of the control group after 24 h. We found significant differences among the groups (***P = 0.00003 × 10^-4^; Fig 9B). A total of 102, 102, and 107 images were obtained for the treatment groups (A and B) and control group (C), respectively. Twenty images from Group A were excluded because of their poor quality. We analyzed 494 measurements from Group A, 613 from Group B, and 641 from Group C.

**Fig 9.**
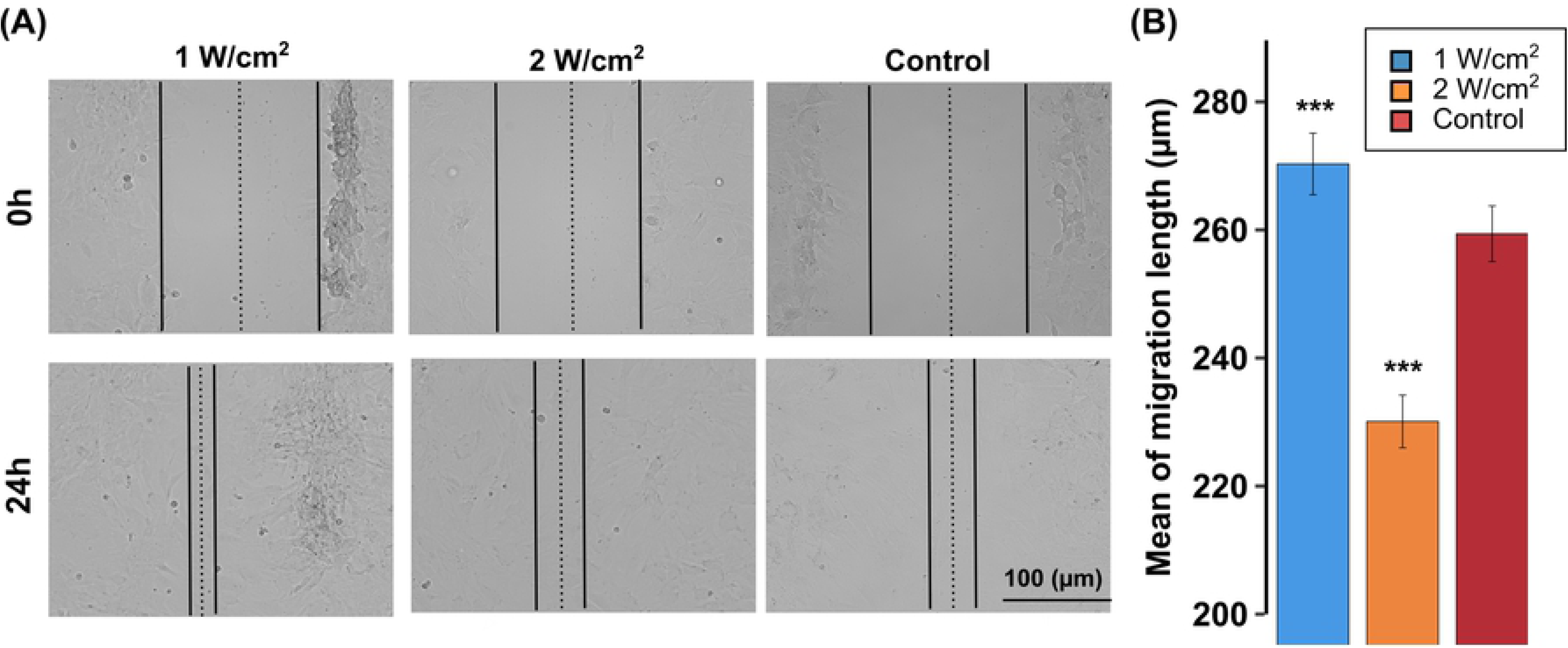
Effect of therapeutic ultrasound on fibroblast cell migration in early treatment. (A) Scratch assay results for the low dose treatment Group A (1.0 W/cm^2^), high dose treatment Group B (2.0 W/cm^2^), and control Group C. (B) Mean migration length (µm) 24 h after treatment. Error bars represent SEMs.

### Effect of therapeutic ultrasound on the OD of ECM: type I collagen, type III collagen, and fibronectin synthesis for early and late treatments

Therapeutic ultrasound significantly increased the median OD among groups of type I collagen (*P = 0.03), type III collagen (*P = 0.02), and fibronectin (**P = 0.003) on the 6^th^ day. Compared to the control group on the 6^th^ day after the stimulation period, the low dose of ultrasound increased the OD of the three proteins: type I collagen by 27%; type III collagen by 59%; and fibronectin by 32%. In contrast, compared to the control group on the 6^th^ day after the stimulation period, the high dose of ultrasound increased the OD of both collagens (type I collagen by 57%; type III collagen by 79%) but decreased fibronectin by 83%. Compared to the control group on the 10^th^ day after the stimulation period, the low dose of ultrasound increased the OD of both collagens (type I collagen by 8%; type III collagen by 37%) but decreased fibronectin by 57%. In contrast, compared to the control group on the 10^th^ day after the stimulation period, the high dose of ultrasound increased the OD of the three proteins: type I collagen by 32%; type III collagen by 51%; and fibronectin by 31% (Fig 10).

**Fig 10.**
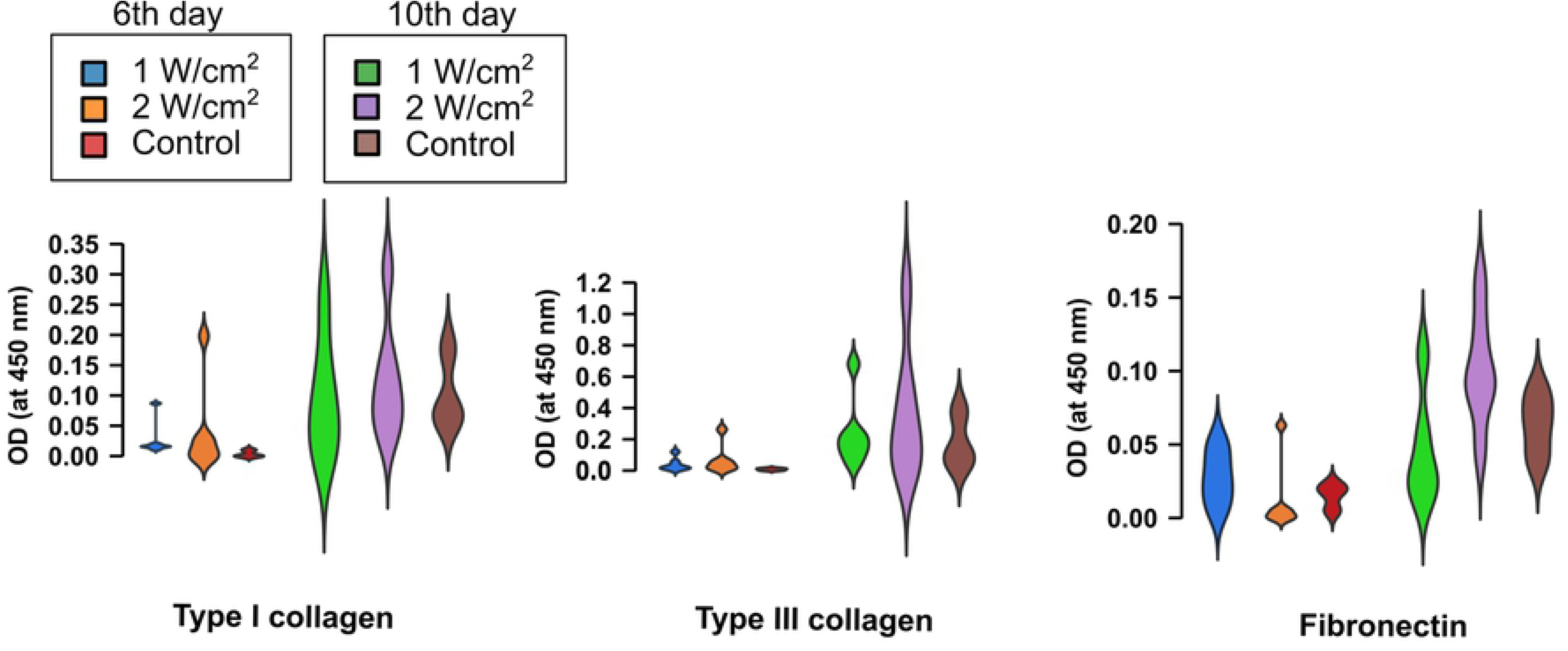
Increase in the optical density (OD) of extracellular matrix (ECM) for early and late treatments due to therapeutic ultrasound. Protein OD at 450 nm of ligament fibroblasts.

Therapeutic ultrasound increased the OD and protein concentration of type I collagen, type III collagen, and fibronectin significantly (**P = 0.005) among the groups on the 6^th^ day. Compared to the control group on the 6^th^ day after the stimulation period, the low dose of ultrasound did not synthesize type I collagen or type III collagen; however, fibronectin was increased by 79%. In contrast, on the 6^th^ day after the stimulation period, the high dose of ultrasound did not synthesize type I collagen or type III collagen, but decreased fibronectin by 100%. Compared to the control group on the 10^th^ day after the stimulation period, the low dose of ultrasound increased the concentration of both collagens (type I collagen by 30%; type III collagen by 33%) but decreased fibronectin by 33%. In contrast, compared to the control group on the 10^th^ day after the stimulation period, the high dose of ultrasound increased the concentration of the three proteins: type I collagen by 45%; type III collagen by 71%; and fibronectin by 44%.

Accordingly, we confirmed that the high dose (2.0 W/cm^2^) of therapeutic ultrasound enhances the synthesis of structural proteins, such as type I collagen, type III collagen, and adhesive proteins such as fibronectin, better than the low dose (1.0 W/cm^2^) on the 10^th^ day after the stimulation period. A box plot shows the protein concentration data (Fig 11).

**Fig 11.**
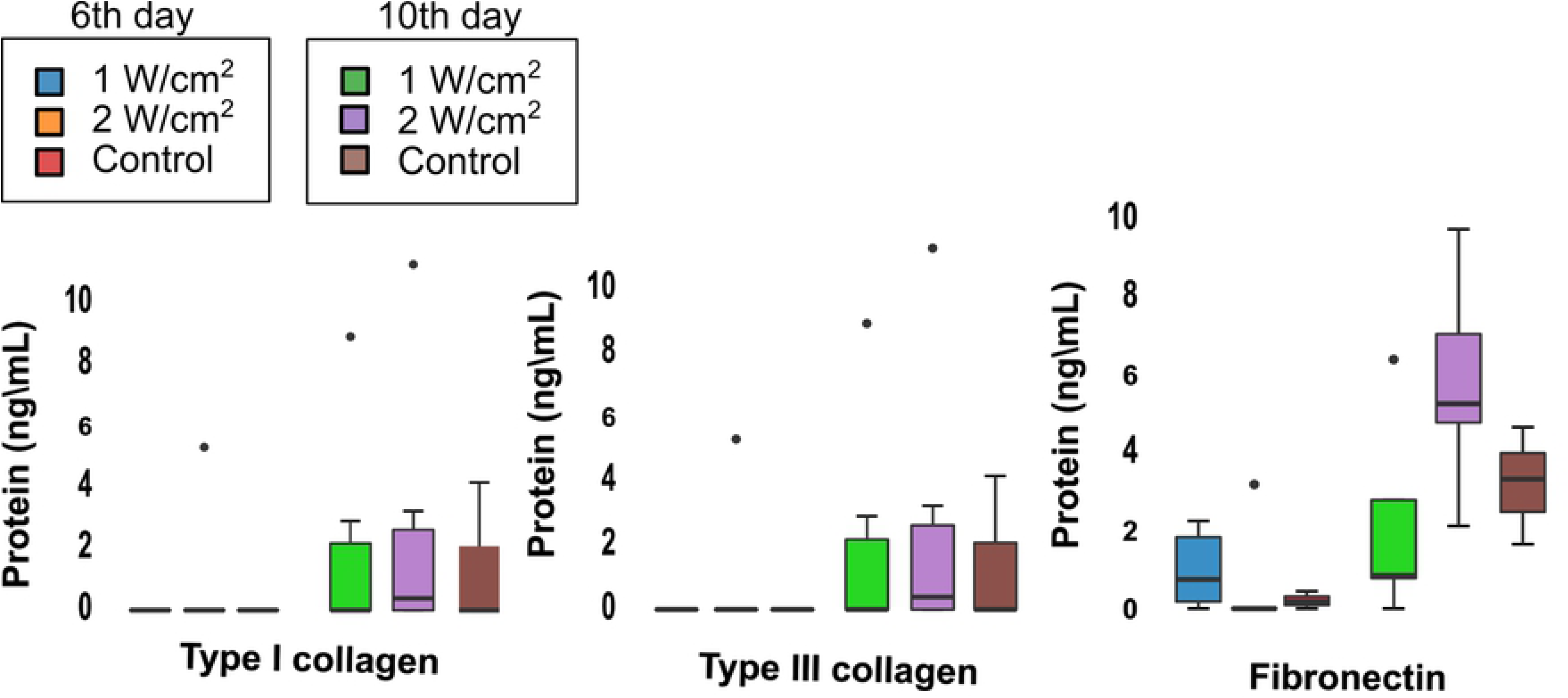
Increase in the concentration of ECM for early and late treatments due to therapeutic ultrasound. Protein concentration in cell culture supernatants of ligament fibroblasts. Values were measured on the 6^th^ day and 10^th^ day after stimulation period for type I collagen, type III collagen, and fibronectin.

### Effect of therapeutic ultrasound on β-actin expression for promoting early and late treatments

Through immunofluorescence imaging, β-actin (an essential structural protein of the cell cytoskeleton) was detected in both treated and control ligament fibroblasts. We found that both doses of therapeutic ultrasound decreased the elastic modulus of ligament fibroblasts, an effect caused by cytoskeleton reorganization, which was visualized for both treatments in our images through β-actin expression (Fig 12). The IHC image analysis showed a greater β-actin shortening area in the treatment groups. Compared to the control group, after the stimulation period, the low dose of ultrasound increased the mean of the β-actin shortening area by 74%. In contrast, compared to the control group, after the stimulation period, the high dose of ultrasound increased the mean of the β-actin shortening area by 31%. Finally, compared to the high dose after the stimulation period, the low dose of ultrasound increased the mean of the β-actin shortening area by 63%. Nevertheless, there was no statistically significant difference among the groups (P = 0.373). This result can be explained by the small sample size for all groups (n=3). Refer to the supporting information (S4 File) for additional details.

**Fig 12.**
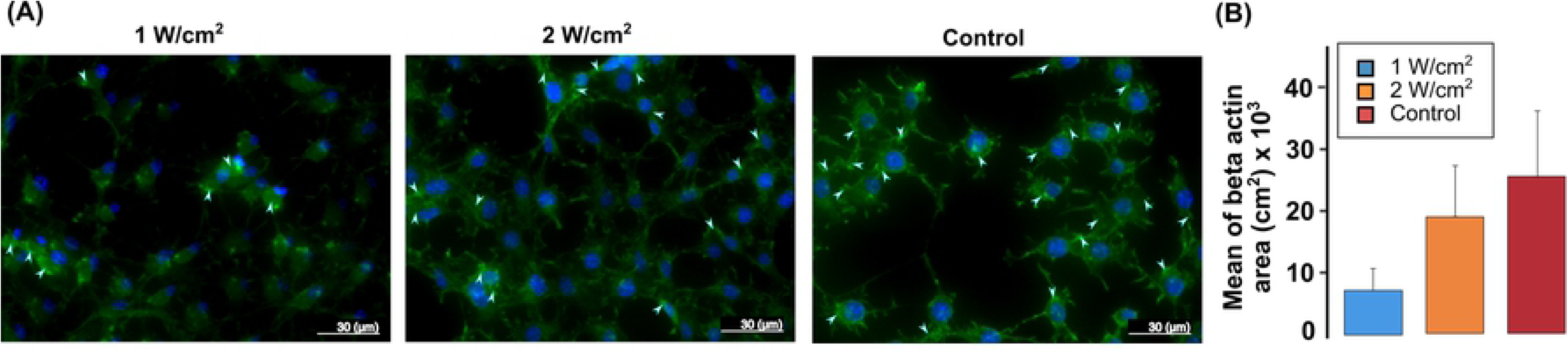
Alteration of β-actin expression to promote early and late treatments due to therapeutic ultrasound. (A) Immunofluorescence analysis of β-actin (green) and cell nuclei (blue) in ligament fibroblasts. (B) Mean of β-actin area (cm^2^). Error bars indicate SEMs.

### Statistical analysis

Descriptive statistics and multiple comparisons between all groups are presented in the supporting information (S4 file).

## Discussion

Through a multidisciplinary approach that included an *in vitro* experiment and a computational simulation, our results demonstrate that conventional doses of therapeutic ultrasound applied for five days modify the mechanical and biological properties of ligament fibroblasts by altering their cytoskeleton while maintaining cell viability. More specifically, low (1.0 W/cm^2^) and high (2.0 W/cm^2^) doses promote cytoskeleton deformation, causing ligament fibroblasts to soften and triggering cell biological responses related to the early (proliferative) and late (remodeling) stages of the wound healing process. Overall, our results suggest that ligament fibroblasts reorganize their cytoskeleton as evidenced by the shortening of the β-actin area [79]. These results are manifested by the decreased elastic modulus, visible cytoskeleton displacements, and decreased harmonics in treated cells [80]. Moreover, both doses activate the dynamic role of the cytoskeleton to (i) allow cells to proliferate and migrate after applying a low dose [81], (ii) increase collagen synthesis to maintain the resistance of the ECM against the high dose [21, 82] and probably restore the fluid volume [79], and (iii) increase the fibronectin synthesis to maintain the cell attached to the surface while collagen is synthesized in the late stage to preserve the shape and cell architecture [21, 82].

The elastic modulus differences between both doses of therapeutic ultrasound and the control group at 1 MHz prove that the resistance required to deform the ligament fibroblast with the AFM tip is lower for the treated groups. As there are no studies on ligament fibroblasts that can be used to compare with our results, in epithelial and endothelial cell lines derived from human breast cancer (MCF-7) and human umbilical vein endothelial cells (HUVEC), low-intensity ultrasound stimulation for 2 s with a frequency of 20 kHz, applied at two intensities of 0.9 and 1.8 W/cm^2^, produced more cell membrane permeability (sonoporation) in HUVEC cells by the cavitation effect of the ultrasound wave, which is attributed to their higher elastic modulus and lower flexibility caused by more organized actin fibers of the cytoskeleton [13].

Another study found a strong relationship between the ultrasound frequency and cell elastic modulus of breast cancer cells. At a frequency of 450 kHz and 60 s exposure to ultrasound, the cell elastic modulus initially tended to increase by 50%. However, when the frequencies were in the range of 550–620 kHz, the cell elastic modulus decreased by 50%. The explanation of this behavior depends not only on the specific interaction between ultrasound doses that produce reorganization of the cytoskeleton but also on the cell density, size, and shape, as well as the ability of cells to detect the stiffness of the neighboring cells and surrounding location [34].

Our findings show that ligament fibroblasts treated with both doses of therapeutic ultrasound tend to be more deformable or flexible but possess the strength to maintain the cell shape without rupture, which can be explained by the disassemblies induced in actin microfilaments [13]. Although greater dispersion is observed, our experimental elastic modulus results (median values) are in the range of the NIH3T3 fibroblasts (0.8–5 kPa) [80, 83]. This dispersion in the elastic modulus may be influenced by the AFM indentation random procedure, which can include one or more of the following: cytoskeleton, membrane, and cell organelles.

In addition, as the cell structure is mainly provided by the cytoskeleton, we inferred that the cell elastic modulus may alter the harmonic vibration of the cell protein network of filaments. Furthermore, while the cytoskeleton filaments had a lower elastic modulus, the natural frequencies of vibration decreased. These results are in agreement with other studies, which affirm that the natural frequencies of vibration of normal cells are higher owing to their high elastic modulus compared to that of tumor cells [7, 84]. On the other hand, the frequency values for the cytoskeleton harmonics differed from the 1 MHz ultrasound frequency. The absence of a resonance effect may explain cell viability. While our goal was to analyze only the cytoskeleton dynamics, the cytoplasm and nucleus were excluded from our simulation. This may explain why the values of the natural frequencies of vibration that we reported (3.3 × 10^9^ to 4.1 × 10^9^ Hz) vary from those obtained in previous studies (21–34 × 10^3^ Hz for tumor cells) [34]. Another reason may be the differences in the elastic modulus of ligament fibroblast cells versus tumor cells [85].

As both ultrasound treatments did not affect the viability of ligament fibroblasts, we confirmed that therapeutic ultrasound does not induce harmful effects by cavitation threshold (bubbles originating in a liquid that interacts with the nearest structure) [37]. In addition, ligament fibroblasts do not perceive the treatments as a negative stimulation because their level of apoptosis is comparable to that of the control group cells. Similar to low-intensity pulsed ultrasound, conventional therapeutic ultrasound at low and high doses is safe and does not affect cell viability or apoptosis [23].

Our results extend the previous findings that fibroblasts treated with ultrasound show increased fibroblast cell proliferation at 1.0 W/cm^2^ [42, 86] and demonstrate that the frequency of ultrasound is not a unique parameter that affects the cell response, as affirmed by Rubin et al. [37]. Our findings imply that external forces produced by different intensities are a plausible reason for the increase in cell proliferation through cytoskeletal reorganization. This means that 1.0 W/cm^2^ of ultrasound stimulates the early stage of wound healing caused by ligament fibroblasts.

Our results are consistent with those of previous studies on other mammalian cells. For example, Tsai et al. demonstrated that ultrasound enhances the migration and proliferation of tendon cells by using doses similar to those in our study [30]. Moreover, Man et al. stated that osteoblast cells exposed to the same frequency as in our study, but with different intensities and time doses, exhibit increased cell migration by 40% [87]. Aterthon et al. demonstrated that low-dose ultrasound treatment enhances the migration speed of osteoblasts (MC3T3) by 30% [88]. Furthermore, Leng et al. showed that low-dose ultrasound treatment increases relative migration by 150% and proliferation by 80% in keratinocyte cells by activating signaling pathways [89]. Although our results have a lower percentage of increased migration than those of other studies, this difference can be explained by the different cell types chosen in the different cited investigations. Thus, we suggest that cell proliferation and migration are dependent on the dosage of ultrasound treatment. These results can be explained by cytoskeleton reorganization, as shown in other studies [23, 88], and by the decrease in the cell elastic modulus of the ligament fibroblasts, as demonstrated in this study.

Our findings regarding collagen synthesis on the 10^th^ day after stimulation are consistent with the observations of Tsai et al., who also found that low intensities of ultrasound (0.1 and 1.0 W/cm^2^) stimulate the synthesis of type I and type III collagen [90]. In terms of fibronectin synthesis, our results for the low dose on the 6^th^ day after stimulation agree with those of Harle et al., who found that fibronectin synthesis was upregulated following stimulation with lower intensities of ultrasound (140 mW/cm^2^) in human osteoblasts. However, for the same low dose, but on the 10^th^ day after stimulation, fibronectin synthesis was diminished as shown with several low intensities of ultrasound (140, 230, 540, and 990 mW/cm^2^) in human periodontal ligament cells [42]. Our results showed that the synthesis of proteins depends not only on the stimulation dose but also on the number of days after stimulation, which indicates the importance of selecting the correct dose of stimulation based on the early or late stage of the ligament healing process. For example, to increase the synthesis of type I and type III collagen in the remodeling or late stage of wound healing, we recommend applying a high dose of stimulation and evaluating the results on the 10^th^ day after stimulation.

In addition, since collagen was not synthesized on the 6^th^ day after stimulation for both the treatment and control groups but was released on the 10^th^ day after stimulation for both doses, we confirmed that collagen production is a complex process of transcription, translation, and assembly to obtain this protein in the ECM [91]. Furthermore, these findings suggest that therapeutic ultrasound may improve the new structural and mechanical ECM of an injured ligament because collagen provides resistance to the tissue [92].

On the other hand, while fibronectin promotes cell adherence to the source surface [93], our results suggest that the low dose on the 10^th^ day after stimulation decreases the synthesis of fibronectin to avoid ligament fibroblast attachment; the decrease in fibronectin enables cell proliferation and migration. A high dose on the 10^th^ day increases the synthesis of fibronectin to enhance ligament fibroblast attachment; the increase in fibronectin facilitates synthesis of type I and type III collagen.

Moreover, we infer that the low dose of ultrasound is the treatment that primarily increased the β-actin shortening area. As shown in living fibroblasts, shortening of stress fibers occurs starting at the proximal end, which is reflected by a decrease in fluorescence intensity as measured in our study [94]. On the other hand, the high dose increased the β-actin shortening area to facilitate collagen and fibronectin synthesis, as we previously demonstrated [95]. To our knowledge, no evidence has been reported about the effects of ultrasound on β-actin expression in ligament fibroblasts; however, low-intensity pulsed ultrasound promotes actin assembly (polymerization) by mechanical stress in osteoblasts [96]. It is also important to mention that a high dose decreases the β-actin shortening area to allow fibroblasts to attach to the ECM and increase collagen synthesis by actin assembly [95].

We focused our attention on the mechanical properties that altered the cytoskeleton dynamics to determine the biological responses related to the wound healing process after applying low and high doses of therapeutic ultrasound on ligament fibroblasts. The elastic modulus and harmonics of the cytoskeleton, such as those observed in cancer cell mechanics, can be used as biomarkers to determine cell function because the elastic modulus is an indicator of cancer cell invasiveness [85], and the harmonics can predict scenarios of possible damage to cells as low-frequency ultrasound induces cytotoxic effects on tumor cells [7]. We propose that the elastic modulus is an indicator of actin reorganization for cell proliferation, migration, and synthesis of collagen and fibronectin after applying therapeutic ultrasound doses to ligament fibroblasts, and that harmonics can be a predictor of cell viability and cytoskeleton deformation. Our findings reveal the reasons for the relevance of the above-described interactions between the cell mechanics and biological responses of ligament fibroblasts caused by the application of therapeutic ultrasound waves.

## Conclusions

Our study introduces a multidisciplinary approach for diagnosing cell function through cell mechanics analysis in ligaments and hopefully in other dense connective tissues such as the tendon, fascia, and skin. This study presents a new perspective of cell mechanics in rehabilitation, which can help researchers engaged in clinical reasoning to formulate specific doses of therapeutic ultrasound that can potentially improve the early and late stages of the ligament wound healing process.

## Author Contributions

The contributions made by each author to the manuscript are presented in Table 3.

**Table 3.**
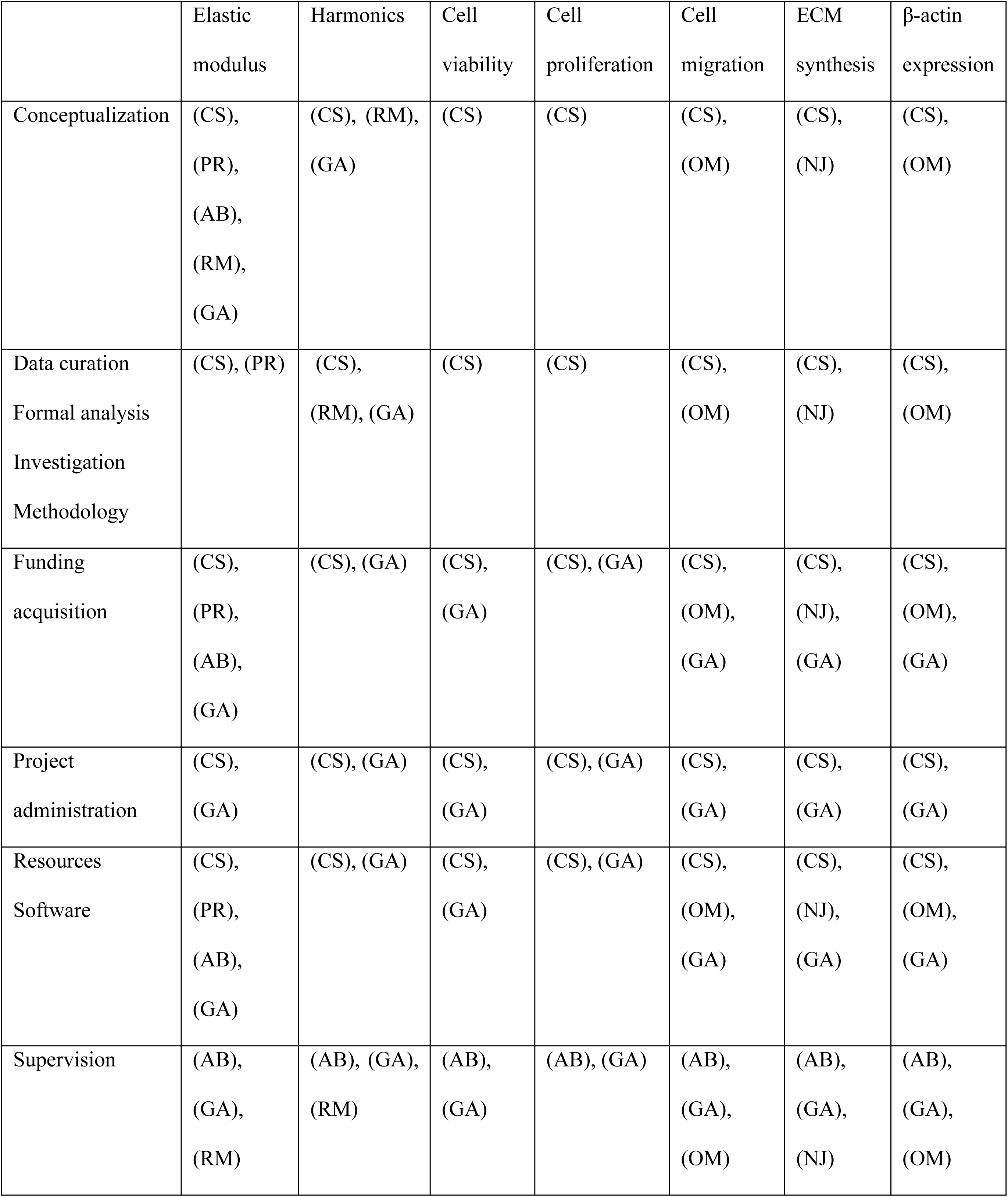
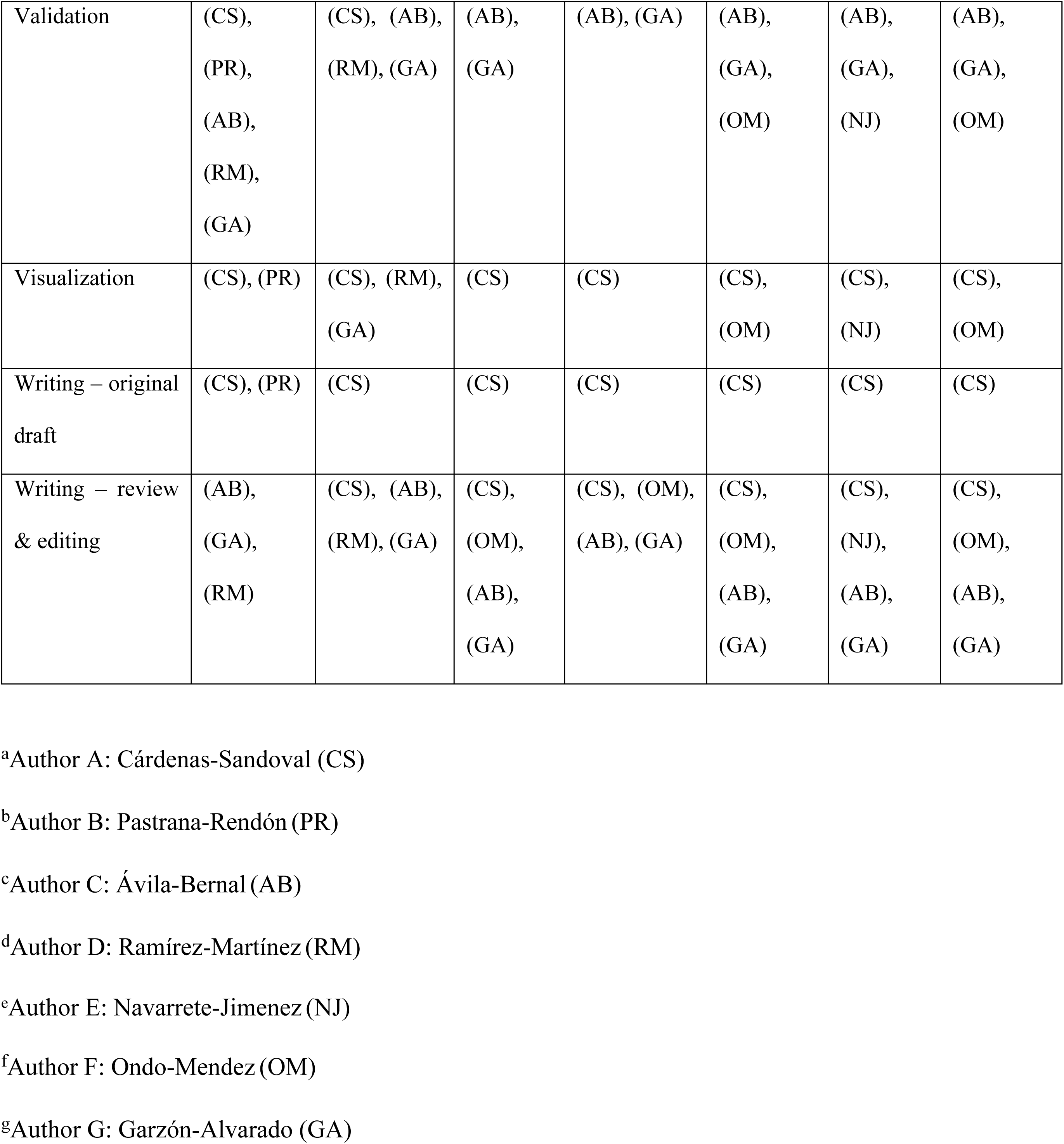
Contributions made by each author to the manuscript.

## Conflict of interest

The authors report no conflicts of interest.

## Supporting information

S1 File: Correction of elastic modulus using the asymptotical correction mode

S2 File: Example of shortening area of β-actin analysis using immunohistochemistry (IHC)

S3 Movies (A-C): Animations of displacements and harmonics of the natural frequency of vibration (50^th^) for all groups

S4 File: Descriptive statistics and multiple comparisons between all groups

S5 File: Python script to create an octahedron tensegrity structure in Abaqus CAE for all groups

## Notes

### Competing Interest Statement

The authors have declared no competing interest.

